# A Region of UNC-89 (obscurin) Lying Between Two Protein Kinase Domains is a Highly Elastic Spring Required for Proper Sarcomere Assembly

**DOI:** 10.1101/2020.04.03.023374

**Authors:** Hiroshi Qadota, Jasmine C. Moody, Leila Lesanpezeshki, Taylor Moncrief, Deborah Kitzler, Purnima Devaki Bhat, Siva A. Vanapalli, Andres F. Oberhauser, Guy M. Benian

## Abstract

In *C. elegans, unc-89* encodes a set of giant multi-domain proteins (up 8,081 residues) localized to the M-lines of muscle sarcomeres and required for normal sarcomere organization and whole-animal locomotion. Multiple UNC-89 isoforms contain two protein kinase domains. There is conservation in arrangement of domains between UNC-89 and its two mammalian homologs, obscurin and SPEG: kinase, a non-domain region of 647-742 residues, Ig domain, Fn3 domain and a second kinase domain. In all three proteins, this non-domain “interkinase region” has low sequence complexity, high proline content and lacks predicted secondary structure. We report that a major portion of this interkinase (571 residues out of 647 residues) when examined by single molecule force spectroscopy *in vitro* displays the properties of a random coil and acts as an entropic spring. We used CRISPR/Cas9 to create nematodes carrying an in-frame deletion of the same 571-residue portion of the interkinase. These animals express normal levels of giant internally deleted UNC-89 proteins, and yet show severe disorganization of all portions of the sarcomere in body wall muscle. Super-resolution microscopy reveals extra, short-A-bands lying close to the outer muscle cell membrane and between normally spaced A-bands. Nematodes with this in-frame deletion show defective locomotion and muscle force generation. We designed our CRISPR-generated in-frame deletion to contain an HA tag at the N-terminus of the large UNC-89 isoforms. This HA tag results in normal organization of body wall muscle, but disorganization of pharyngeal muscle, small body size, and reduced muscle force, likely due to poor nutritional uptake.

**Highlights:**

- The giant muscle proteins UNC-89 and its mammalian homologs have an ∼700 aa non-domain region lying between two protein kinase domains
- By single molecule force spectroscopy UNC-89 non-domain region is an elastic random coil
- Nematodes lacking this non-domain region have disorganized sarcomeres and reduced whole animal locomotion
- UNC-89 non-domain region is required for proper assembly of A-bands from thick filaments

## Introduction

*C. elegans* UNC-89 is the founding member of the UNC-89/obscurin family of giant muscle proteins. Loss-of-function mutations in the *C. elegans* gene *unc-89* result in slow moving worms with disorganized muscle sarcomeres, including a lack of M-lines [1-3]. *unc-89* is a complex gene: through the use of three promoters and alternative splicing at least 16 polypeptides are generated, ranging in size from 156,000 to 900,000 Da [3-5]. Overlapping sets of isoforms are expressed in body wall muscle, pharyngeal muscle (for pumping in and grinding up bacterial food), egg laying and intestinal muscles [3], and several gonadal epithelial cells [6]. The largest of these isoforms, UNC-89-B and UNC-89-F, consist of 53 Ig domains, two Fn3 domains, a triplet of SH3, DH and PH domains near their N-termini, and two protein kinase domains (PK1 and PK2) near their C termini. Antibodies localize UNC-89 to the M-line [3,4]. To learn how UNC-89 is localized and performs its functions, our lab has systematically identified its binding partners. We have screened a yeast two hybrid library with segments representing the largest UNC-89 isoform and identified at least 7 interacting partners, and 6/7 have orthologs or homologs in humans [7-13]. Interactions have been verified by a combination of demonstrating binding in vitro using purified recombinant proteins, co-immunolocalization, and identifying muscle mutant phenotypes.

Mammalian striated muscle express three UNC-89-like proteins from separate genes, obscurin (OBSC), obscurin-like 1 (obsl1) and striated muscle preferentially expressed gene (SPEG). UNC-89 is most similar to obscurin [14,15], which also expresses many isoforms, including giant isoforms (obscurin A), and shorter isoforms with 2 protein kinase domains [16]. Although UNC-89 is located only at the M-line, various obscurin isoforms are located at either the M-line, or the Z-disk [15,17]. Mutations in the human OBSC gene result in cardiomyopathies (HCM, DCM, LVNC)[18]. In addition to serving as a platform for assembly of M-line proteins, obscurin links myofibrils to the sarcoplasmic reticulum (SR). The C-terminus of obscurin interacts with an ankyrin localized to the SR [19,20]. The obscurin KO mouse has normal myofibril organization but disorganized SR [21]. This myofibril to SR linkage is conserved in nematode UNC-89 [22]. Obsl1 is an ∼200 kDa protein consisting solely of 20 tandem Ig domains, and in heart muscle is localized to M-lines, Z-disks and intercalated disks [23]. Several portions interact with titin and myomesin at the M-line [24]. However, obsl1 is ubiquitously expressed, global KO is embryonic lethal and mutations in the human Obsl1 gene result in a growth defect (3M-growth syndrome)[25]. SPEG also exists as different sized isoforms, but the largest isoform is 3,267 aa and consists of 9 Ig and 2 Fn3 domains, and 2 protein kinase domains [26]. It can be regarded as similar to the C-terminal tandem kinase domain regions of obscurin and UNC-89. SPEG is located in a complex lying between the SR and plasma membrane (JMC) in cardiomyocytes. In the mouse, cardiac-specific KO results in DCM, whereas mutations in human SPEG result in centronuclear myopathy with DCM [26]. The *in vivo* substrates for the kinase domains of UNC-89, obscurin and SPEG are not known. However, sequence analysis suggests that: (i) UNC-89 PK1 is catalytically inactive, whereas PK2 is an active kinase [27]; (ii) human obscurin PK1 and PK2 are both likely to be active kinases [27], and each has been shown to undergo autophosphorylation *in vitro* [28]. Obscurin PK2 can phosphorylate N-cadherin in vitro [28]. There is a suggestion that SPEG phosphorylates the JMC protein junctophilin 2 [29].

As shown in Fig. 1, UNC-89, obscurin and SPEG have an overall conservation in organization around the kinase domains: PK1, a non-domain region of 647-742 residues, Ig, Fn, and PK2. The non-domain interkinase region has low sequence complexity, high proline content (11.8—13.3%) and without predicted α-helix and β-strands, are likely to be random coils [9]. These properties of the non-domain interkinase region, henceforth referred to as interkinase (IK), are shared with the known highly elastic spring regions of human titin (PEVK and N2A/B)[30]. In this paper, we describe, for nematode UNC-89, the response of the interkinase to mechanical pulling and the phenotype of worms in which the interkinase has been deleted in-frame. This is the first time the function of this region has been explored for any member of the UNC-89/obscurin family.

**Fig. 1.**
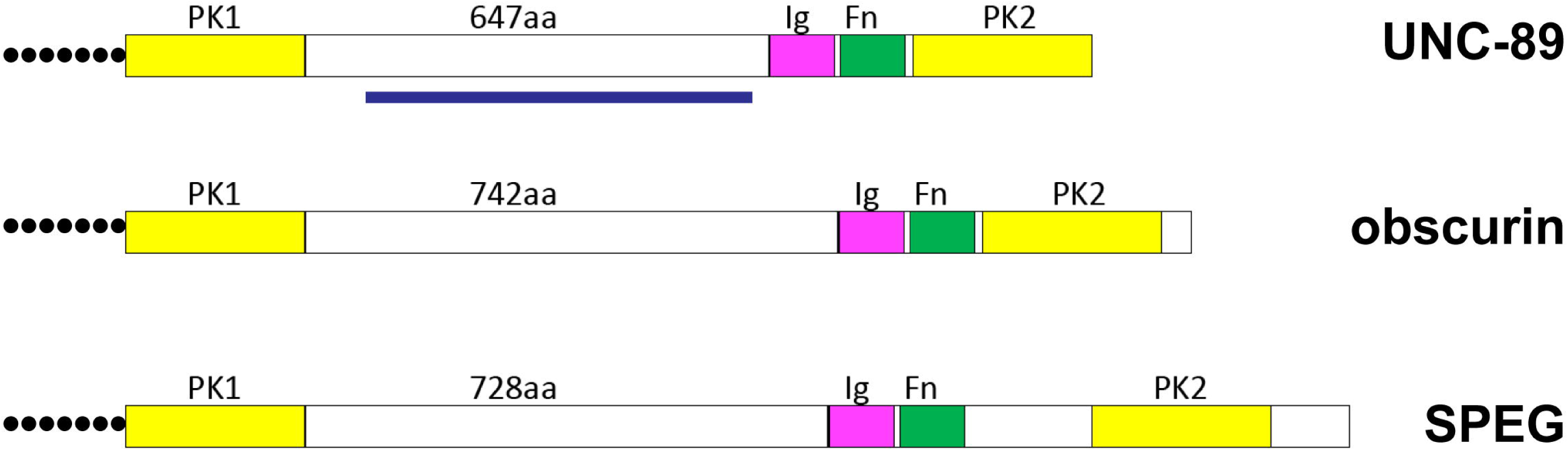
Schematic representation of domain organization around kinase domains for nematode UNC-89, and mouse obscurin and SPEG. Note that each protein, near its C-terminus consists of a protein kinase, a non-domain region (647-742 residues), Ig and Fn domains, and a second protein kinase domain.

## Results

Given the nature of the amino acid sequence of the interkinase region of UNC-89, we wished to determine how it might respond to a mechanical pulling force using the single molecule force spectroscopy technique. To carry this out, we first expressed in E. coli 1 571 residue segment comprising the majority of the 645 residue interkinase region (IK) flanked on the N- and C-terminus by two copies of Ig27 from human titin. The response of Ig27 to pulling force is well documented and served as an internal control. In addition, the protein had a 6XHis tag at its N-terminus (Fig. 2A). After purification (Fig. 2B), the protein was allowed to attach to a glass surface coated with Ni^2+^-NTA to allow binding of the N-terminus of this polyprotein via its 6XHis tag. The stylus of the single-molecule force spectrometer was allowed to bind non-specifically and at random points along the polyprotein, and then pulling force was applied by moving the piezo-electric platform downward. A representative force curve of the mechanical unfolding of the polyprotein in which the stylus had attached to the C-terminus is shown in Fig. 2C. After a long and featureless trace without energy barriers, the typical unwinding of the 4 Ig domains is found. The statistical distribution of the contour length of the long featureless peak (Fig. 2D) shows that the mean contour is approximately 185 nm. This is a good match to the expected contour length of a fully unwound polypeptide of 571 amino acids, since the average length of an amino acid is 0.34 nm, i.e. 0.34 nm/aa X 571 aa = 194 nm. Based on this force curve and contour length, we can conclude that the IK has the mechanical properties of a random coil and behaves like an entropic spring.

**Fig. 2.**
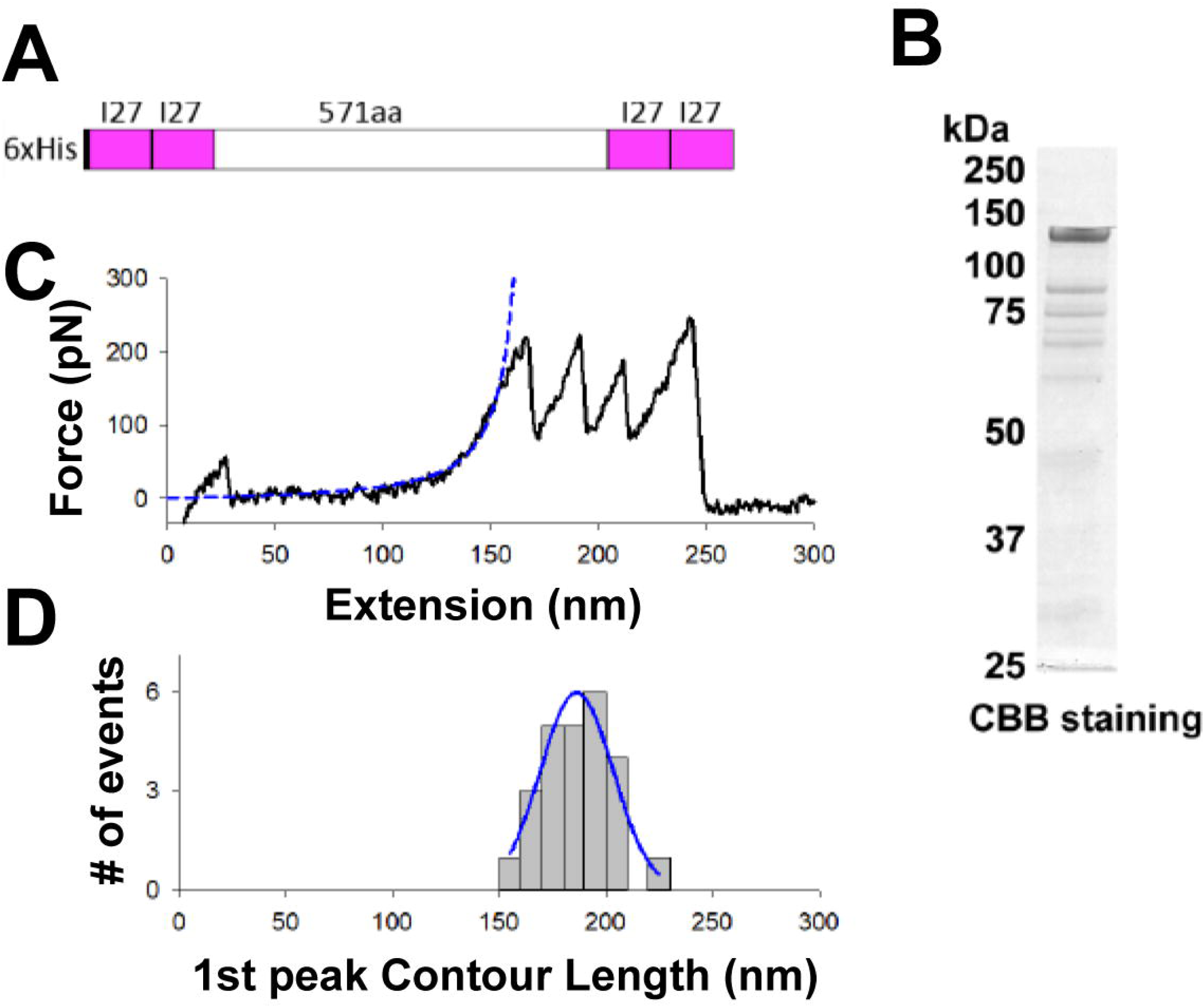
The interkinase region (IK) of UNC-89 is a highly elastic spring. (**A**) Polyprotein construct encompassing the IK flanked on either side by two titin I27 domains used in single molecule force spectroscopy (SMFS) experiments. (**B**) SDS-PAGE after Coomassie Blue staining of ∼2 μg of the polyprotein shown in (A). (**C**) Representative force curve of the mechanical unfolding of the polyprotein by SMFS. The dotted line is a worm-like-chain (WLC) fit to the force-extension curve with free contour length L_C_ and a fixed persistence length L_p_ = 0.36 nm. (**D**) Statistical distribution of the contour length of the first peak (L_C_ = 186 ± 3 nm, n=18). The solid line represents a Gaussian fit of the histogram.

We wished to determine the *in vivo* consequences of removing the IK from UNC-89. To do this, we used CRISPR/Cas9 to create an in-frame deletion of the same 571 residues that we explored with the single molecule method in vitro. However, to be able to monitor the expression of UNC-89 large isoforms in which the IK had been deleted we added an HA tag to the N-terminus of the large isoforms. We thought such tagged UNC-89 could also be used in future immunoprecipitation experiments to identify proteins in a complex with UNC-89. Animals expressing HA-UNC-89 large isoforms were obtained by CRISPR/Cas9, and the strain is designated *unc-89(syb797)*. As shown later, this HA tag does not affect the localization of UNC-89 in body wall muscle, nor does it affect the organization of body wall muscle sarcomeres, or whole animal locomotion. However, although we were able to detect HA-UNC-89 in body wall muscle by immunostaining (data not shown), we were unable to detect HA-UNC-89 by western blot. Perhaps the HA epitope is buried in a portion of the UNC-89 N-terminus. Nevertheless, we used *unc-89(syb797)* as starting point to make an in-frame deletion of 571 residues of the IK, and this strain is designated *unc-89(syb797 syb1257)*. A schematic of the genomic region deleted is shown in Fig. 3a. Before characterizing either strain further, each was outcrossed 5X to wild type, to remove most possible off-target effects of CRISPR/Cas9 gene editing.

**Fig. 3.**
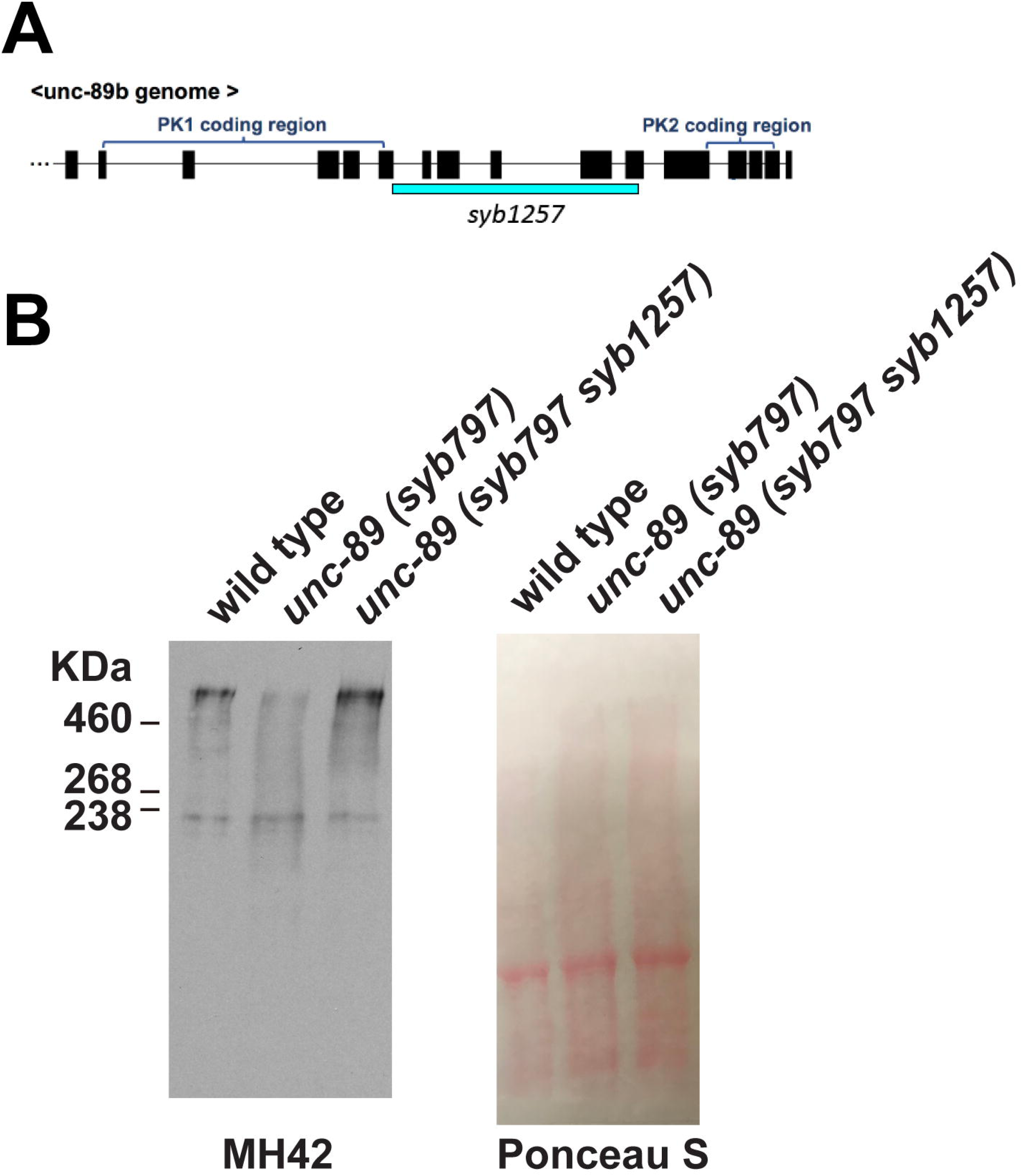
*unc-89(syb797 syb1257)* is an in-frame deletion of most of the IK region and does not affect the level of expression of large UNC-89 isoforms. (A) Representation of the exon (black boxes) and intron (lines) organization of the kinase-encoding region of the unc-89 gene. Blue bar, portion deleted in syb1257 by CRISPR/Cas9. This is a 1,713 bp in-frame deletion which internally deletes 571 aa of the IK beginning 66 aa C-terminal of PK1 and continuing through 10 aa N-terminal of Ig53. (B) Western blot demonstrating that the large UNC-89 isoforms are expressed in *unc-89(syb797)* and *unc-89(syb797 syb1257)*. Total Laemmli-soluble proteins from the indicated strains were separated on a 5% SDS-PAGE, transferred to membrane and reacted with a monoclonal antibody to UNC-89 (MH42). On the right is shown Ponceau S staining of the blot before reaction with antibodies. The band at∼230 kDa likely represents a degradation product.

As shown in Fig. 3B, neither addition of the HA tag (in *syb797*), nor the in-frame deletion of IK (in *syb797 syb1257*), leads to appreciable decrease in the levels of large UNC-89 isoforms by western blot. We had hoped to be able to use an antibody to PK2 to prove that the deletion is in-frame, however, two attempts to make such an antibody (in rabbits or rats) failed. Nevertheless, since the levels of large isoforms did not diminish, it is likely that we achieved an in-frame deletion which does not destabilize the protein.

To assess the *in vivo* consequences of removing the IK, we examined the organization of body wall muscle sarcomeres, and whole animal locomotion and force generation. To assess sarcomere organization, we determined the immunolocalization of several proteins with known sarcomere locations using confocal microscopy; F-actin for thin filaments, myosin heavy chain A for thick filaments, UNC-89 for M-lines, UNC-95 for the bases of M-lines and dense bodies (Z-disk analogs), and ATN-1 (α-actinin) for the deeper portion of dense bodies. As shown in Fig. 4 (left), the HA tag (in *syb797*) does not appreciably affect the organization of sarcomeres as the patterns observed appear identical to that of wild type muscle. However, deletion of IK (in mutant *syb797 syb1257*) results in disorganization of sarcomeres in all substructures (Fig. 4 right). It is important to note however that in *syb797 syb1257*, although the pattern of UNC-89 is disorganized (i.e. not parallel straight lines as in *syb797* or wild type), UNC-89 is still detectable at high levels consistent with the western blot result. Note also that in *syb797 syb1257*, the disorganization of UNC-89 appears quite similar to the pattern of disorganization of myosin heavy chain A (MHC A). This is not unexpected as thick filaments in nematode body wall muscle have differential localization of two myosin heavy chain isoforms, with MHC A in the middle, and MHC B in the major outer portions [31]. Normally thick filaments are arranged parallel and in register, so that the M-line is the structure in the middle of the A-band, which contains MHC A, and where the thick filaments are crosslinked. Thus, when imaging *C. elegans* body wall muscle with light microscopy, the localization of MHC A and UNC-89 are nearly identical [4,32]. Note that the bases of M-lines are also wavy and discontinuous upon anti-UNC-95 staining (Fig. 4, right inset). The disorganization in UNC-95 localization is so severe that often rows of dense bodies overlap with M-lines. Overall, these results indicate that the IK is required for the proper organization of sarcomeres.

**Fig. 4.**
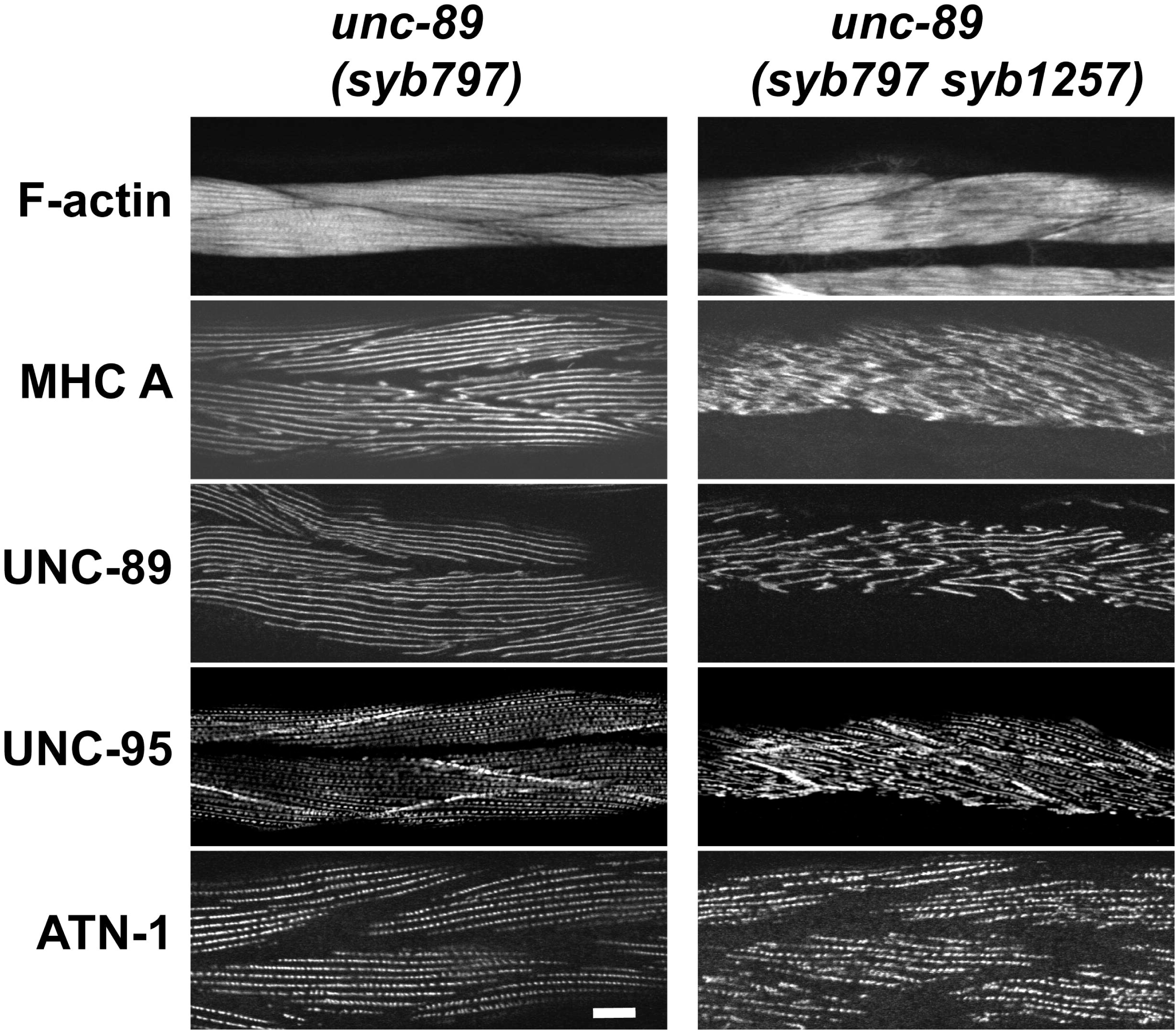
*unc-89(syb797 syb1257)* animals that express HA-UNC-89 isoforms with an in-frame deletion of the interkinase region have disorganized myofibrils, whereas *unc-89(syb797)* animals that express HA-UNC-89 isoforms have normally organized myofibrils. Each panel shows representative confocal images of several body wall muscle cells stained with antibodies to the indicated sarcomeric components or with rhodamine-phalloidin. Scale bar, 10 μm.

We examined the organization of MHC A and UNC-89, proxies for the middle of A-bands, at higher resolution using structured illumination microscopy (SIM). In Fig. 5A and B, it is clear that the N-terminal HA tag on the large UNC-89 isoforms (in *syb797*) has no effect on the normal pattern of straight parallel A-bands. However, in-frame deletion of IK (in *syb797 syb1257*), for both MHC A and UNC-89 immunostaining, it is evident that the A-bands are broken, and much of the parallel arrangement is missing. In addition, there appear to be extra, short A-bands (indicated by arrows in Fig. 5) that lie in the spaces between otherwise nearly normal A-bands. By capturing optical sections with an interval of 0.2 μm, we noticed that these partial A-bands lie mostly close to the outer cell membrane of the muscle cells. From top to bottom of Fig. 5A and B, images are shown from close to the outer muscle cell membrane to deeper into the cell. Note that the partial A-bands lie near the outer muscle cell but do not extend as deeply as normal A-bands.

**Fig. 5.**
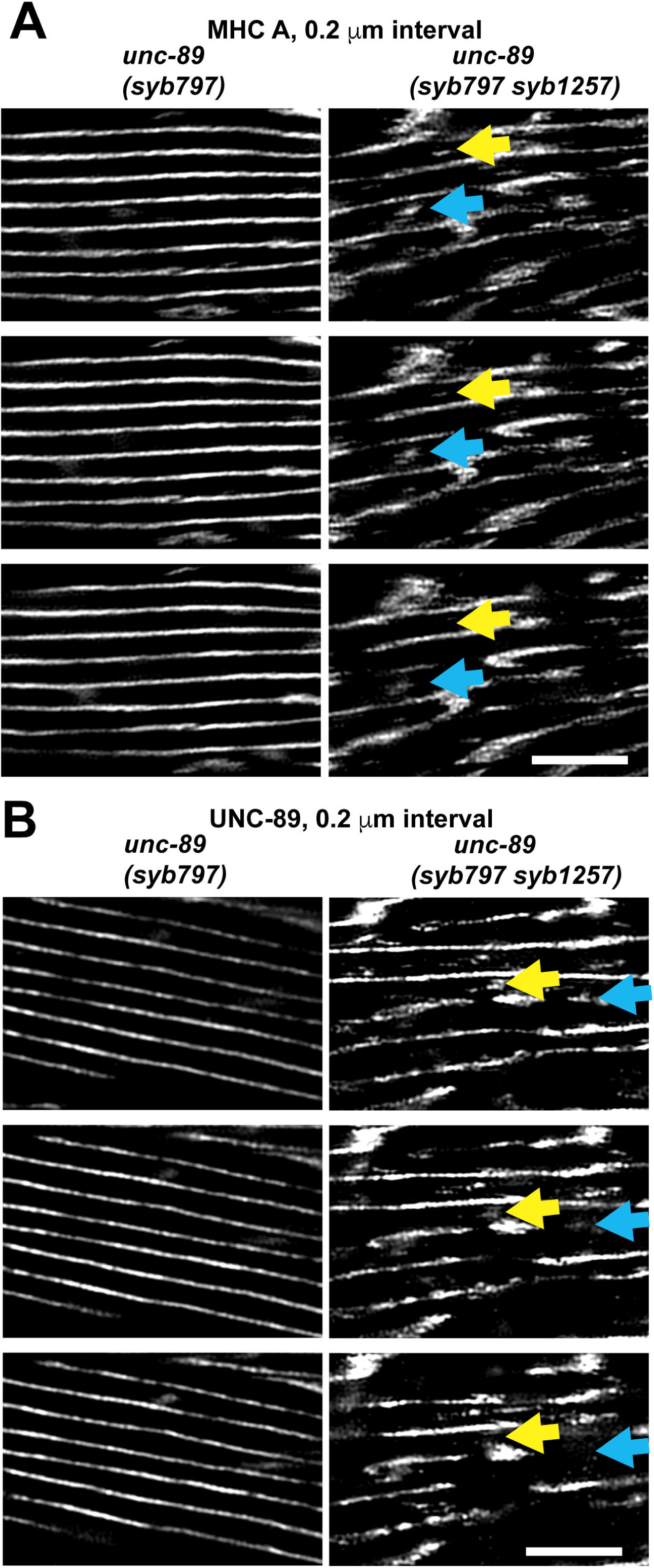
*unc-89(syb797 syb1257)* animals that express HA-UNC-89 isoforms with an in-frame deletion of the interkinase region show extra A-bands that are not full-depth. After immunostaining with antibodies to MHC A (A) or antibodies to UNC-89 (B) multiple optical slices with a 0.2 μm interval were obtained by structured illumination microscopy (SIM). Yellow and blue arrows point to examples of short extra A-bands that lie between normally located A-bands and reside close to the outer muscle cell membrane. In comparison, *unc-89(syb797)* that expresses HA-UNC-89 shows parallel, normally spaced and full-depth A-bands. Scale bar, 5 μm.

In *C. elegans*, the body wall muscle is required for the animal to generate the forces required for its normal swimming, crawling and burrowing locomotion [33]. Given the disorganization of sarcomeres in the body wall muscle of nematodes expressing large isoforms of UNC-89 with an in-frame deletion of the IK, we wondered if these animals would display defects in movement or in muscle force generation. First, because our strain with the IK deletion also contains an N-terminal HA tag, we measured locomotion in the strain with the HA tag alone, *syb797*. As shown in Fig. 6A and B, *syb797* is not slower in swimming or crawling as compared to wild type. In fact, *syb797* displays a statistically faster swimming than wild type. However, clearly, *syb797 syb1257*, containing both an HA tag and IK deletion shows a 15% reduction in swimming and a 30% reduction in crawling (Fig. 6A and B). This decrease in swimming speed is comparable to that displayed by the canonical *unc-89* allele *e1460*, but not as severe as that of alleles *su75* or *tm752* (Fig. 6A). The decrease in crawling of the IK deletion strain is not as severe as 3 of 4 other alleles tested (*e1460, su75, tm752*). In terms of muscle force exerted, as measured by the NemaFlex assay [34], all mutant alleles (including *syb797 syb1257*) except for *tm752* are defective compared to wild type (Fig. 6C).

**Fig. 6.**
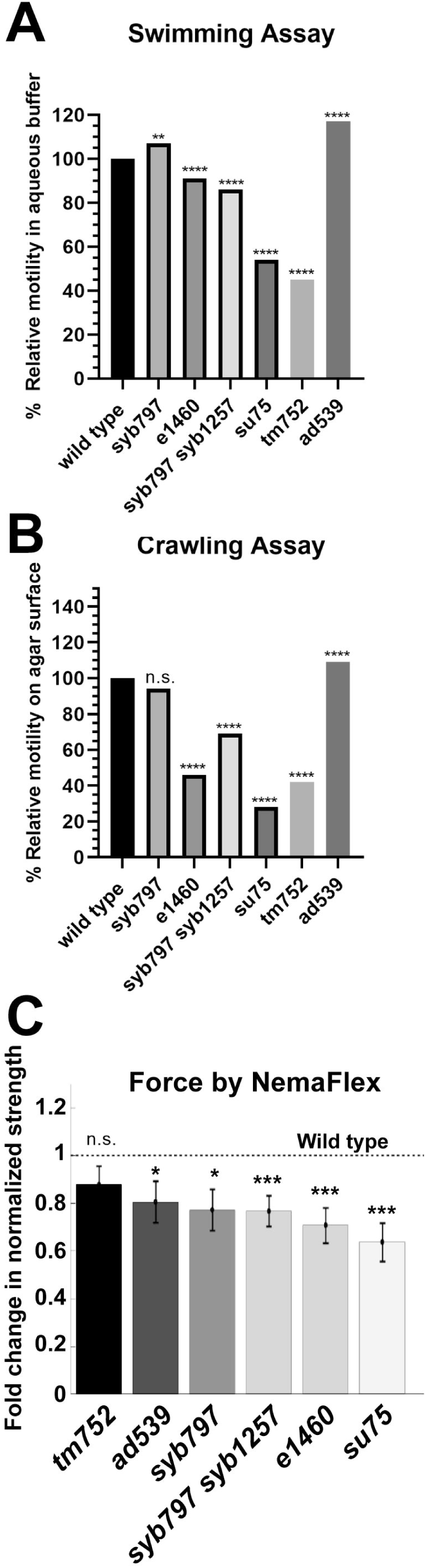
Whole worm locomotion and muscle force measurements of wild type and various *unc-89* mutant alleles. (A) Swimming assays show that *unc-89(syb797)* and *unc-89(ad539)* swim faster than wild type, whereas all other *unc-89* mutants, including *unc-89(syb797 syb1257)* swim slower than wild type. (B) Crawling assays show that *unc-89(ad539)* crawls faster than wild type, that *unc-89(syb797)* crawls at the same speed as wild type, and all other *unc-89* mutants crawl more slowly than wild type. C) NemaFlex force measurements show that *unc-89(tm752)* develops the same muscle force as wild type, whereas all other *unc-89* mutants develop less force than wild type, although, *unc-89(syb797 syb1257), unc-89(e1460)*, and *unc-89(su75)* are more severely affected. The actual data, including sample sizes, means, and standard errors are shown in Supplementary Table 1 for swimming and crawling and Supplementary Table 2 for NemaFlex measurements.

Interestingly, *syb797*, which has an HA tag at the N-terminus of the large isoforms, shows no decrease in crawling, but an increase in swimming, and decreased ability to generate force. This phenotype of increased speed of swimming, no change or even increased speed of crawling, and yet decreased force, is shared with the allele *ad539. unc-89(ad539)*, like *syb797*, has normally organized sarcomeres in body wall muscle, which likely accounts for lack of any decrease in swimming or crawling. (We first showed that *unc-89(ad539)* has normally organized sarcomeres by EM [2], but to verify this result we have immunostained its muscle with a variety of antibodies to sarcomeric proteins, and all of these are normally localized (Supplementary Figure 1)).

In *C. elegans*, the pharynx is a neuromuscular pump required for bringing bacterial food into the worm and grinding it up before its entry into the intestine. *unc-89(ad539)* was first isolated in a screen for mutants that show defective feeding behavior, and found to be an allele of *unc-89* by complementation testing and 3 factor mapping; by polarized light microscopy the pharyngeal muscle was shown to have reduced birefringence and abnormal arcs in the terminal or posterior bulb [35]. Based on this, and the similarity of *ad539* with *syb797*, we examined the pharyngeal muscle structure of both these alleles, and additional alleles of *unc-89* using polarized light microscopy (Fig. 7). As first noted by Waterston et al. [1], in *unc-89(e1460)*, instead of the radially oriented filaments observed in the posterior bulb of wild type, this mutant has longitudinally oriented filaments at the periphery of the bulb (indicated by yellow arrow in Fig. 7). In fact, all the *unc-89* mutants that we imaged show this defect except for *tm752*, which shows normal pharyngeal muscle organization. Remarkably, *syb797*, with the HA tag, shows the same defect. However, *ad539* shows not only the defect in the terminal bulb; it also shows a defect in the anterior bulb (blue arrows in Fig. 7). The only other *unc-89* allele that shows both bulbs affected is *su75*.

**Fig. 7.**
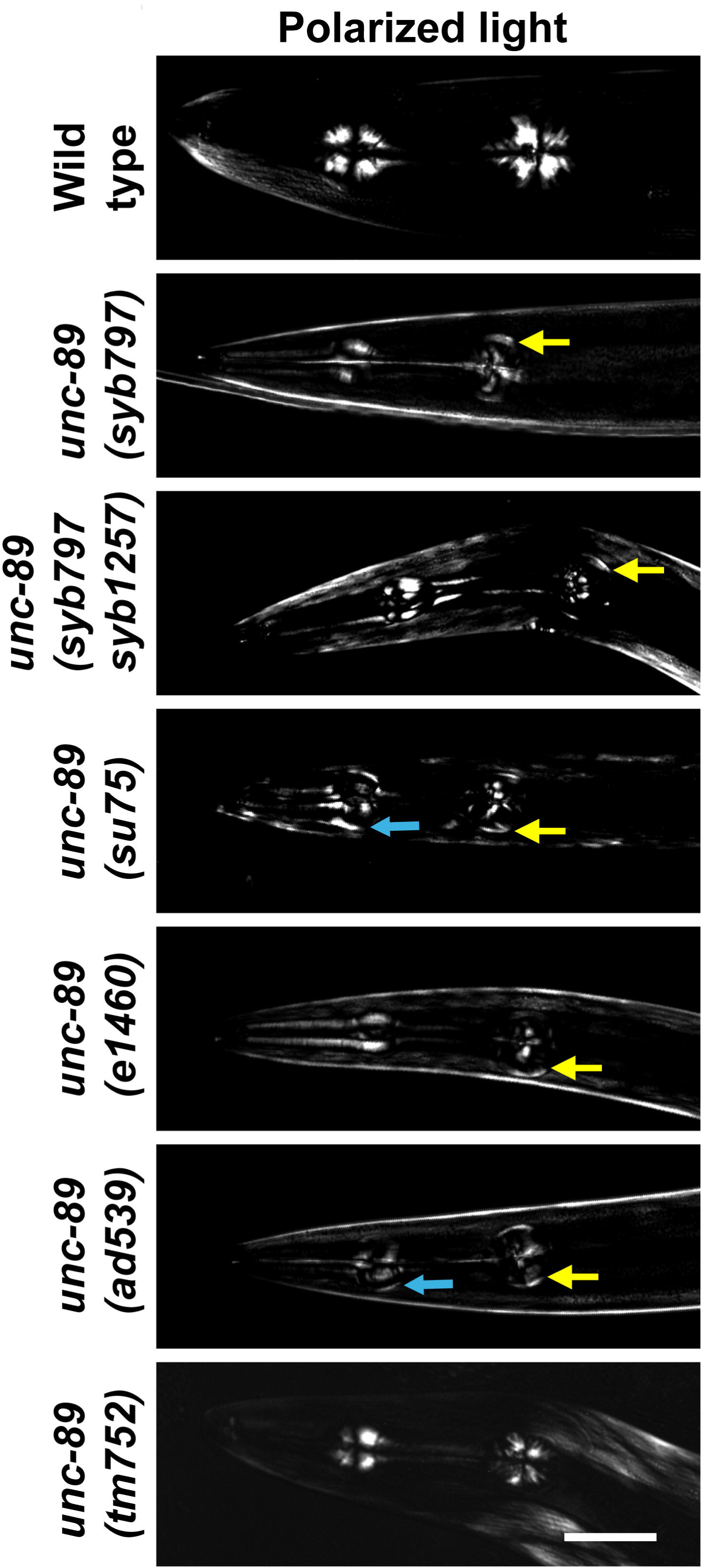
Polarized light images of the myofilament lattice in the pharynges of wild type and various *unc-89* mutant alleles. Yellow arrows indicate mis-localized arc-like thick filaments at the periphery of the terminal bulb, and blue arrows indicate mis-localized arc-like thick filaments at the periphery of the anterior bulb. Note that *unc-89(tm752)* shows pharyngeal muscle like wild type, whereas all the other *unc-89* mutants show defective organization. *unc-89(su75)* and *unc-89(ad539)* are the only alleles that show defects in both the posterior and anterior bulbs. Scale bar, 50 μm.

To understand the pharyngeal muscle defect further, we examined the expression of UNC-89 protein in the pharynges of wild type, *syb797* and *ad539*. This was done by co-immunostaining worms with antibodies to UNC-89 and to the pharyngeal myosin heavy chain MHC C [36]. We have previously demonstrated that in wild type, UNC-89 is localized to the middle of thick filaments in pharyngeal muscle (Benian et al. 1996). As shown in Fig. 8, UNC-89 is localized to the middle of thick filaments throughout the pharynx. However, in *syb797* there is no detectable expression of UNC-89 in the posterior bulb. In *ad539* there is no observable expression of UNC-89 throughout the pharynx. Moreover, MHC C staining reveals that in both *syb797* and *ad539*, there are abnormal accumulations of thick filaments at the periphery of the posterior bulb (indicated by yellow arrows in Fig. 8), corresponding to the arcs observed by polarized light. The expression defects in the mutants seem to correspond with the defects in pharyngeal muscle organization and indicate that UNC-89 is required for normal organization of sarcomeres in pharyngeal muscle.

**Fig. 8.**
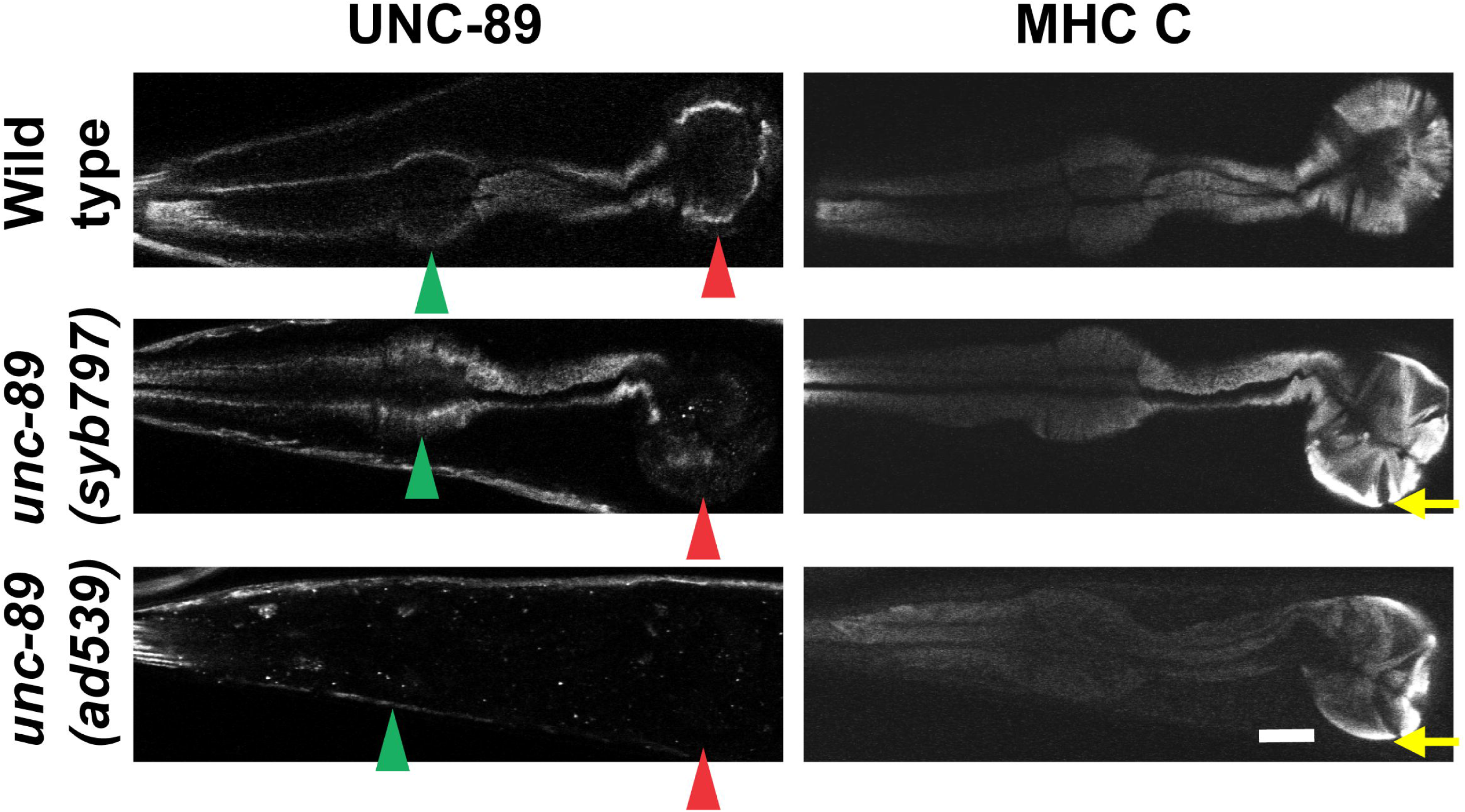
*unc-89(syb797)* and *unc-89(ad539)* have reduced levels of expression of UNC-89. The indicated strains were immunostained with antibodies to UNC-89 and the pharyngeal-specific myosin heavy chain C (MHC C), and representative confocal images of the pharynx are shown. Note that *unc-89(syb797)* shows reduced UNC-89 expression in the terminal bulb, whereas *unc-89(ad539)* shows reduced UNC-89 expression in both the terminal and anterior bulbs. Green arrowhead, anterior bulb; red arrowhead, posterior bulb; yellow arrow, arc-like thick filaments mis-placed at the periphery of the posterior bulb. Scale bar, 10 μm.

## Discussion

In this study, we explored the biophysical properties and *in vivo* function of an example of ∼600 aa sequence that has similar features (low sequence complexity, lack of predicted domains and even secondary structure elements) and location between two protein kinase domain in the UNC-89/obscurin class of giant muscle proteins. By use of single molecule force spectroscopy *in vitro* we showed that this sequence, which we call IK, is a random coil and acts like an entropic spring. The nature of this sequence (random coil) and its physical properties (entropic spring) is shared with the highly elastic spring elements (PEVK, N2B) of another giant muscle protein, titin. In the case of titin, there is good evidence to suggest that these elements unwind when muscle sarcomeres are stretched beyond their relaxed state, store potential energy and then provide a passive restoring force to allow the sarcomeres to return to their resting (relaxed) lengths [37].

To explore the muscle function of the IK, we used CRISPR/Cas9 to create a worm strain, *syb797 syb1257*, in which most of the IK of UNC-89 was deleted in-frame. Despite expressing approximately normal levels of internally deleted UNC-89 isoforms, these animals displayed highly disorganized sarcomeres in their body wall muscle cells. The normal location of UNC-89 is the M-line, the structure in which thick filaments are cross linked in the middle of the A-band. Deletion of the IK, in addition to disrupting the normal continuous and parallel M-lines or A-bands, results in formation of short A-bands which are not full-depth but confined to the portion nearest the outer muscle cell membrane, and these occur between normally spaced A-bands. Sarcomere assembly is known to occur from the outside inwards, first with the deposition of ECM proteins like perlecan, followed by localization of integrins, and then localization of integrin adhesion proteins (UNC-112 (kindlin), PAT-4 (ILK), UNC-97 (PINCH), PAT-6 (α-parvin), followed by assembly of M-line or dense body (Z-disk)-specific proteins [38,39]. We know that UNC-89 interacts with CPNA-1 [11], which in turn interacts with PAT-6, a member of the 4-protein complex that interacts with the cytoplasmic tail of PAT-3 (β-integrin)[40]. It should be noted that CPNA-1 interacts with Ig1-Ig3, which is near the N-terminus of UNC-89 and remote from IK. Our results seem to indicate that these protein interactions at the membrane are not sufficient for full assembly of a M-line/A-band, and the proper spacing between A-bands. Either the length or the elasticity of UNC-89 IK is required. In future experiments, we can determine the minimum length of IK that is sufficient for full A-band assembly, and whether elasticity is crucial. We could replace the elastic IK with a sequence of similar length that is rigid. An example of a rigid sequence that could be used is the series of 4 consecutive Ig domains in human titin that are not separated by linker sequences [41]. It is tempting to speculate that the length of the interkinase is important for somehow determining the spacing between adjacent A-bands. Another possibility is that deletion of the interkinase in some way interferes with the activity of the active protein kinase domain of UNC-89 PK2. This is not likely because inactivation of PK2 does not result in a defect in sarcomere organization (Matsunaga et al., manuscript in preparation).

In vertebrate muscle, the M-band is known to be elastic and helps titin maintain the registration of adjacent thick filaments during muscle contraction [42]. The cross-linking of thick filaments occurs by the binding of antiparallel dimers of myomesin to myosin of thick filaments. The M-band also includes complex interactions between Obsl1 and obscurin with titin, and myomesin with obscurin [43]. Although much of the elasticity and mechanical protection is provided by myomesin [44,45], our results suggest that the IK region of obscurin, which also lies in the M-band also provides some of this elasticity.

Because whole animal locomotion (swimming, crawling) and force generation are diminished upon removal of the IK, we also have learned that the level of sarcomere organization found in this mutant is not sufficient for proper contractile activity. In contrast, the better level of sarcomere organization in *unc-89*(*tm752)* is sufficient to generate normal force but is not sufficient to generate normal swimming or crawling speeds.

Our IK deletion strain was also contains an HA tag appended to the N-terminus of the large UNC-89 isoforms. We designed this for future immunoprecipitation experiments to further test whether UNC-89 interacting partners that we have discovered primarily through two hybrid screens are actually associated with UNC-89: we would create in-frame deletions of segments of UNC-89 that are suspected to bind to a partner, then IP those internally deleted UNC-89 proteins, and look for absence of the association. Although the HA does not appreciably affect the level of expression of large UNC-89 isoforms, and does not affect the organization of sarcomeres in body wall muscle or the ability of whole animals to swim or crawl, it does reduce the ability to generate muscle force. In addition, the HA tag results in a defect in thick filaments in the terminal bulb of the pharynx. Moreover the level of expression of UNC-89 is reduced in the terminal bulb. This suggests that the HA tag at the N-terminus either interferes with transcriptional regulation of pharyngeal-specific UNC-89 isoforms, or interferes with interaction with binding partners found in pharyngeal muscle. Although our extensive yeast two-hybrid screens did not reveal any binding partners for the N-terminus of UNC-89, these screens cannot reveal all relevant binding partners. The reduced force generated by *syb797* containing the HA tag is also puzzling. However, it should be noted that *syb797* and *ad539* share these properties—each has normal body wall muscle, but defective pharyngeal muscle. Then, why do these two mutants generate less force? In fact, the only mutant studied that generates normal force is *tm752* that does so despite having mildly disorganized body wall muscle. The key to explaining why these two mutants generate less force appears to be their reduced body diameters. In fact, as shown in Table 1, all the mutants generating less force have smaller than wild type body diameters, and defective pharynxes. *tm752* has a normal body diameter and a normal pharynx. Our speculation is that all the mutants with defective pharynxes have reduced nutrition and consequently are smaller. Although our force measurements were normalized for body diameter, it is possible that reduced nutrition results in reduced metabolism and reduced energy production, and thus less ability to generate force. It is also curious that both *syb797* and *ad539* display greater swimming speeds and *ad539* shows greater crawling speed. Perhaps this is because these thinner animals can bend more easily.

**Table 1:** The effects of *unc-89* mutations on expression of UNC-89 isoforms, body wall and pharyngeal muscle structure, locomotion, body wall muscle force and body diameter.

| Strain | Mutation | Affect on Expression | Body Wall Muscle Structure | Swimming | Crawling | Force by NemaFlex | Body Diameter (μm) | Pharyngeal Muscle |
| --- | --- | --- | --- | --- | --- | --- | --- | --- |
| wild type |  |  |  |  |  |  | 73.5–78.3 |  |
| *syb797* | HA tag at N-term. of large isoforms | reduced in pharynx | normal | faster | normal | reduced | 62.1 | defective posterior bulb |
| *e1460* | nonsense mutation in alternatively spliced exon for Ig20–Ig26 [22] | reduced levels of large isoforms [12] | moderate [12] | slower | slower (severe) | reduced (severe) | 57.3 | defective posterior bulb |
| *syb797 syb1257* | HA tag added to N-term. of large isoforms & in-frame deletion of IK | normal expression of large isoforms | moderate–severe | slower | slower | reduced | 55.7 | defective posterior bulb |
| *su75* | 1 bp insertion in coding for Ig38 resulting in frameshift premature stop [12] | all giant isoforms missing small kinase isoforms present [3,12] | severe [12] | slower (severe) | slower (severe) | reduced (severe) | 49.9 | defective posterior & anterior bulbs |
| *tm752* | out of frame deletion in IK [5] | all kinase isoforms (giant & small) missing [5] | mild [12] | slower (severe) | slower (severe) | normal | 75.9 | normal |
| *ad539* | Ala to Val change in KSP region but this may not be causative | normal body wall no detectable in pharynx | normal | faster | faster | reduced | 56.2 | defective posterior & anterior bulbs |

## Materials and Methods

### Recombinant protein production

A cDNA encoding a 571 amino acid portion of the interkinase region of UNC-89, IYDYLRIQ…ADIEKVTP (residues 6945—7515 in UNC-89b) was synthesized using E. coli optimal codons and cloned into pUC57 by GenScript USA,. Inc. (Piscataway, New Jersey). This plasmid was digested with SpeI and SacI, and the insert was cloned into “I27 plasmid shuffled codons (003)”, in order to express a fusion protein consisting of 6xHis-I27-I27-UNC-89 IK 571 aa—I27-I27. (I27 is the human titin Ig27 domain.) The resulting plasmid was transformed into E. coli BL21 (DE3) and the fusion protein was expressed at 20° for 5 hours after induction with 750 mM Isopropyl β-D-1-thiogalactopyranoside (IPTG). The 6xHis tagged fusion protein was purified with Ni-NTA Agarose (Qiagen), as described in [13]. The buffer of the eluted protein was exchanged with PBS containing 10% glycerol.

### Single-molecule force spectroscopy (SMFS) experiments

Pulling experiments using SMFS were carried out on a home-built single-molecule Atomic Force Spectrometer as previously described in the literature [46,47]. In each experiment, we deposited 10 μL of the UNC-89 IK domain polyprotein solution (0.5 mg/mL) in PBS (pH 7.4) onto a Ni^2+^-NTA functionalized glass coverslip [48] covered by 50 μL of PBS. We allowed the protein to absorb onto the substrate for 10 minutes before pulling. The spring constant of each individual cantilever (MLCT, Bruker AFM Probes, Camarillo, CA) was determined experimentally by measuring the cantilever power spectrum and using the equipartition theorem [49,50] at the beginning of each experiment. All measurements were done at a constant velocity of 500 nm/s in PBS solution at room temperature. A worm-like chain (WLC) model [51] with a variable persistence length was fit to the first I27 peak to measure the contour-length (Lc) of the IK domain in the force extension data [52].

### Generation of CRISPR/Cas9 edited nematodes

Strains PHX797 and PHX1257 were generated by SunyBiotech (http://www.sunybiotech.com) using CRISPR/Cas9 technology in the following ways. PHX797, *unc-89(syb797)*, was created from wild type strain N2 to contain an HA tag at the N-terminus of the large UNC-89 isoforms, using sgRNAs with the following sequences: sg1: ccatcATGGCTAGTCGACGCCAAAAG, and sg2: cccaccttacccttaccatcATGG. PHX1257 *unc-89(syb797 syb1257)* was created from PHX797 to contain a 4,614 bp in-frame deletion of the interkinase region, using sgRNAs with the following sequences: Sg1:CCTGTAGCACCG**G**AAGGTCGACG, Sg2:CCACCACCGACTGTGGAATACGT, Sg3:AACAGCAATTGAGAGAATTAAGG and Sg4:CCACAAAGAAGAATGATGATGGA.

Both strains were outcrossed 5 times to wild type to remove most of the possible off target effects of CRISPR/Cas9; PHX797 to GB312, and PHX1257 to GB313.

### Antibodies and western blots

We used the procedure of Hannak et al. [53] to prepare total protein lysates from wild type, *unc-89(syb797)* and *unc-89(syb797 syb1257)* animals. Equal amounts of total protein from these strains were separated by a 5% separating and 3% stacking Laemmli SDS-PAGE gel, transferred to nitrocellulose membrane for 2 hours and reacted with anti-UNC-89 MH42 mouse monoclonal [4,54] at 1:100 dilution, followed by reaction with anti-mouse Ig-horseradish peroxidase, and ECL.

### Immunofluorescence microscopy

Adult worms were fixed as described previously [55,56]. For body wall muscle straining, these primary antibodies were used at the following dilutions: anti-myosin heavy chain A (MHC A; mouse monoclonal 5-6; [31]) at 1:200; anti-UNC-89 [4] (rabbit polyclonal EU30) at 1:100; anti-UNC-95 (rabbit polyclonal Benian-13; [57]) at 1:200; and anti-ATN-1 (α-actinin; MH35; [58]) at 1:200. For pharyngeal muscle staining, the fixed worms were incubated with anti-UNC-89 (rabbit polyclonal EU30; [4]) at 1:100, and anti-myosin heavy chain C (MHC C; mouse monoclonal 9.2.1; [36]), at 1:50. MH35 antibodies were kindly provided by Dr. Pamela Hoppe (Western Michigan University). 5-6 and 9.2.1 antibodies were obtained from the Developmental Studies Hybridoma Bank, created by the NICHD of the NIH and maintained at The University of Iowa, Department of Biology, Iowa City, IA 52242. The secondary antibodies included anti-mouse IgG-Alexa 594 (Invitrogen A11032), and anti-rabbit IgG Alexa 488 (Invitrogen A21206), each at 1:200 dilution. Images shown in Figures 4 and 8, and the Supplemental Figure were captured at room temperature with a Zeiss confocal system (LSM510) equipped with an Axiovert 100M microscope and an Apochromat ×63/1.4 numerical aperture oil objective. The color balances of the images were adjusted by using Adobe Photoshop. We examined the staining of at least 3 worms in each strain.

Super-resolution microscopy was performed with a Nikon N-SIM system in 3D structured illumination mode on an Eclipse Ti-E microscope equipped with a 100×/1.49 NA oil immersion objective, 488- and 561-nm solid-state lasers, and an EM-CCD camera (DU-897, Andor Technology). The color balances of the images were adjusted by using Adobe Photoshop (Adobe, San Jose, CA). A Z-series at 0.2 μm intervals was taken from the outer muscle cell membrane deeper into the muscle cell, as shown in Figure 5.

### Nematode locomotion assays

#### Swimming Assay

Day 2 adults from two, 6 cm NGM OP50 seeded plates were washed off the plates, washed free from bacteria and collected into M9 buffer such that the ratio of worms to buffer was 1:1. Two ml of M9 buffer was added to an unseeded 6 cm NGM plate where upon 5 μl of worm suspension was added to the center of the plate. Worms were allowed 5 minutes to adapt before a video recording of their swimming motions was made using a dissecting stereoscopic microscope fitted with a CMOS camera (Thorlabs). Ten, 10 second videos were recorded for each nematode strain from different sections of the plate, each video tracking the motion of ∼10 worms. The video data was analyzed by Image J FIJI WrmTracker software [59] to obtain body bends per second (BBPS) from a total of 48-83 individual worms for each strain. The resulting worm tracks for each animal were observed, with immobilized animals in addition to upper and lower range outliers being removed from BBPS values for each strain prior to the statistical analysis. The final values for each strain were compared to wild type and tested for statistical significance using a Student T-test.

#### Crawling Assay

Day 2 adults were harvested as above, except that all washing steps used M9 buffer containing 0.2 g/L gelatin. 5 μl of worm suspension was added to the center of a 6 cm unseeded NGM plate, and the excess liquid was removed. After 5 minutes for adaptation, worm crawling was recorded using the above-mentioned strategy for extraction of body bends per second (BBPS) from a total of 66-90 individual worms for each strain. The resulting values for each strain were compared to wild type and tested for statistical significance using a Student T-test.

### Measurement of *C. elegans* muscle force using NemaFlex

The maximum exertable muscle force by *C. elegans* strains was measured using the NemaFlex technique as previously described [34]. The technique is based on deflection of soft micropillars as the animals are crawling through the micropillar arena. The micropillar devices had pillars arranged in a square lattice with a pillar diameter of 44 μm and height of 87 μm. The gap between the pillars is 71 μm. Age synchronized day 1 adult animals were loaded individually in each chamber [60], and a 1-minute video was captured for each animal in a food-free environment at a temperature of 20 ± 1 °C. Imaging was performed in brightfield using a Nikon Ti-E microscope and Andor Zyla sCMOS 5.5 camera. Images were acquired at 5 frames per second with a 4x objective at a pixel resolution of 1.63 μm per pixel. Movies were then processed and analyzed for force values using our in-house-built image processing software (MATLAB, R2016a). Animal force measurements were calculated by identifying the maximal force exerted in each frame for an individual animal. We define the maximum exertable force *f*_95_ corresponding to the 95^th^ percentile of these maximal forces. To account for differences in animal body diameter, we normalize *f*_95_ by the cube of animal body diameter [34]. To compare muscle forces of mutants with wild-type animals, we take the ratio of this normalized maximum exertable force and denote it as fold change in muscle force. Statistical analysis was performed using Wilcoxon rank-sum test.

### Polarized light microscopy

Polarized light images were captured with a Zeiss Axioskop microscope equipped with 20X/0.40 Pol strain-free objective and an AxioCam MRm digital camera and software (Carl Zeiss, Jena, Germany).

## Glossary

## Abbreviations

Ig domain: immunoglobulin domain;
Fn3 domain: fibronectin type 3 domain;
SPEG: striated muscle preferentially expressed gene;
HA: hemagglutinin

## Acknowledgements

This study was supported in-part by a grant from the National Institutes of Health (R01 AR064307) to G.M.B. J.C.M. was supported in-part by the National Science Foundation-Graduate Research Fellowship Program (Grant #DGE 1444932). T.M. was partially supported by a grant from the National Institutes of Health (R01GM118534) to A.F.O. and G.M.B. L.L. and S.A.V. were partially supported by the National Aeronautics and Space Administration (Grant # NNX15AL16G).

---

**Supplementary Fig. 1.**
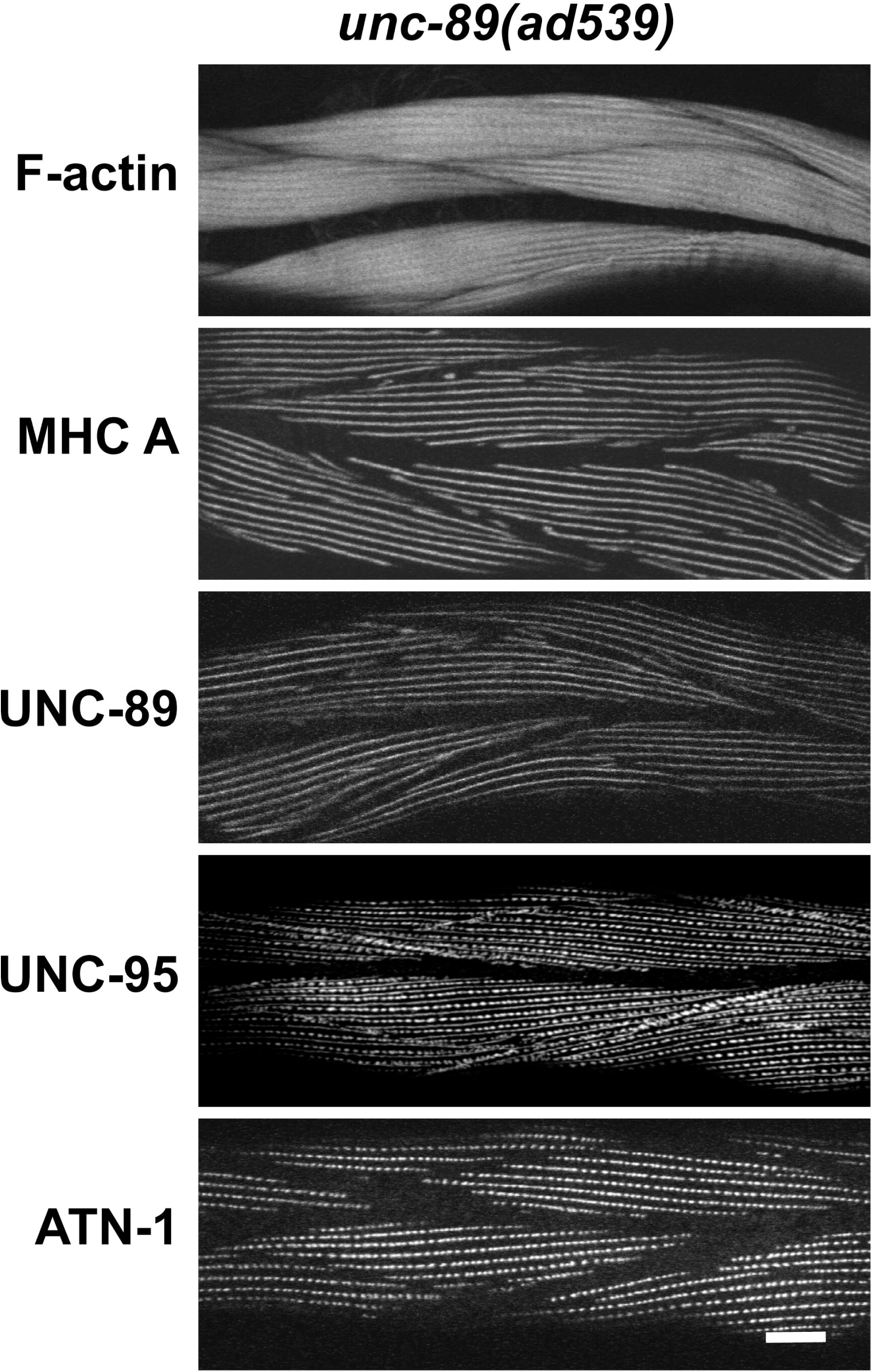
The body wall muscle of *unc-89(ad539)* has normally organized sarcomeres. Each panel shows representative confocal images of several body wall muscle cells stained with antibodies to the indicated sarcomeric components or with rhodamine-phalloidin for F-actin. Scale bar, 10 μm.

**Supplementary Table 1.**
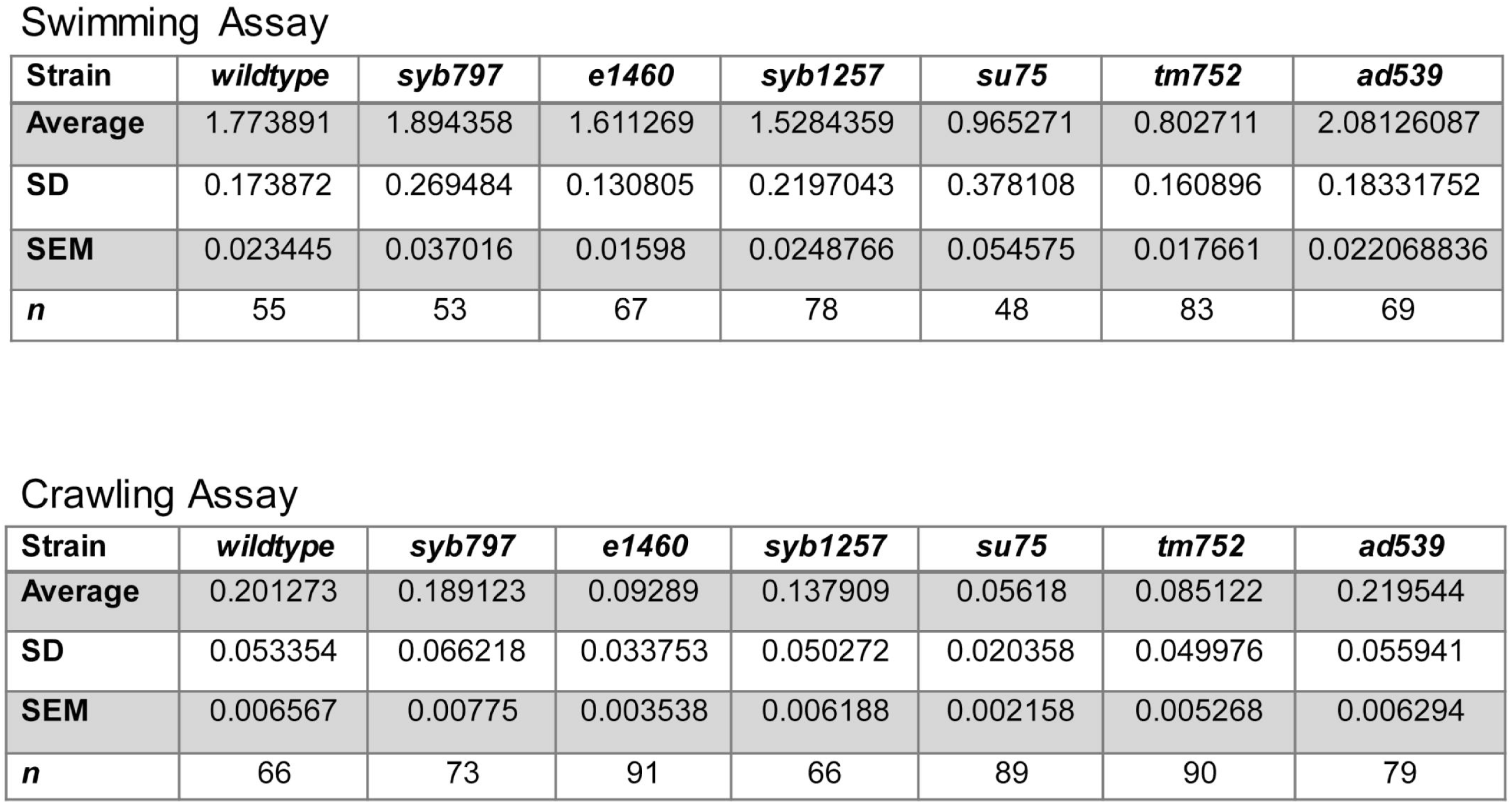
Detailed statistics for swimming and crawling assays on the different strains used in this study and summarized in Fig. 6A and B.

**Supplementary Table 2:** Muscle force and body diameter data for the different strains used in this study and summarized in Fig. 6C.

| Mutant | Sample size | $f_{95}$ | | Normalized $f_{95}$ | | Body diameter | | Confinement* | |
| --- | --- | --- | --- | --- | --- | --- | --- | --- | --- |
| | | Mean ($\mu\text{N}$) | SEM | Mean($\mu\text{N}/\mu\text{m}^3$) | SEM | Mean ($\mu\text{m}$) | SEM | Mean | SEM |
| WT | 36 | 31.3 | 1.88 | $6.710^{-5}$ | 0.37 | 77.3 | 0.38 | 1.09 | 0.006 |
| *unc-89(tm752)* | 31 | 26.3 | 1.97 | $5.910^{-5}$ | 0.35 | 75.9 | 0.56 | 1.06 | 0.009 |
| WT | 30 | 31.3 | 1.86 | $6.510^{-5}$ | 0.35 | 78.3 | 0.39 | 1.10 | 0.006 |
| *unc-89(e1460)* | 33 | 8.6 | 0.35 | $4.610^{-5}$ | 0.14 | 57.2 | 0.54 | 0.80 | 0.008 |
| *unc-89(su75)* | 33 | 5.1 | 0.25 | $4.110^{-5}$ | 0.20 | 49.9 | 0.45 | 0.70 | 0.007 |
| WT | 17 | 28.8 | 1.73 | $7.210^{-5}$ | 0.40 | 73.7 | 0.54 | 1.04 | 0.007 |
| *unc-89(syb797)* | 17 | 13.3 | 0.91 | $5.510^{-5}$ | 0.36 | 62.1 | 0.37 | 0.88 | 0.006 |
| *unc-89(syb797 syb1257)* | 20 | 9.5 | 0.42 | $5.510^{-5}$ | 0.18 | 55.7 | 0.60 | 0.78 | 0.008 |
| WT | 20 | 26.6 | 1.97 | $6.710^{-5}$ | 0.44 | 73.4 | 0.37 | 1.04 | 0.006 |
| *unc-89(ad539)* | 26 | 9.5 | 0.53 | $5.410^{-5}$ | 0.29 | 56.2 | 0.52 | 0.79 | 0.008 |

\*Confinement is defined as the ratio of animal body diameter at mid-length to the gap between pillars.

## References

[1] R.H. Waterston, J.N. Thomson, S. Brenner, Mutants with altered muscle structure in C. elegans, Dev. Biol. 77 (1980) 271–302.

[2] G.M. Benian, A. Ayme-Southgate, T.L. Tinley, The genetics and molecular biology of the titin/connectin-like proteins of invertebrates, Rev. Physio. Biochem. Pharm. 138 (1999) 235–268.

[3] T.M. Small, K.M. Gernert, D.B. Flaherty, K.B. Mercer, M. Borodovsky, G.M. Benian, Three new isoforms of C. elegans UNC-89 containing MLCK-like protein kinase domains, J. Mol. Biol. 342 (2004) 91–108.

[4] G.M. Benian, T.L. Tinley, X. Tang, M. Borodovsky, The Caenorhabditis elegans gene unc-89, required for muscle M-line assembly, encodes a giant modular protein composed of Ig and signal transduction domains, J. Cell Biol. 132 (1996) 835–848.

[5] T.M. Ferrara, D.B. Flaherty, G.M. Benian, Titin / connectin-related proteins in C. elegans: a review and new findings, J. Musc. Res. Cell Motil. 26 (2005) 435–447.

[6] K. Ono, R. Yu, S. Ono, Structural components of the nonstriated contractile apparatuses in the C. elegans gonadal myoepithelial sheath and their essential roles for ovulation, Dev. Dyn. (2007) 236, 1093–1105.

[7] H. Qadota, L.A. McGaha, K.B. Mercer, T.J. Stark, T.M. Ferrara, G.M. Benian, A novel protein phosphatase is a binding partner for the protein kinase domains of UNC-89 (obscurin) in C. elegans, Mol. Biol. Cell 19 (2008a) 2424–2432.

[8] H. Qadota, A. Blangy, G. Xiong, G.M. Benian, The DH-PH region of the giant protein UNC-89 activates RHO-1 GTPase in C. elegans body wall muscle, J. Mol. Biol. 383 (2008b) 747–752.

[9] G. Xiong, H. Qadota, K.B. Mercer, L.A. McGaha, A.F. Oberhauser, G.M. Benian, A LIM-9 (FHL) / SCPL-1 (SCP) complex interacts with the C-terminal protein kinase regions of UNC-89 (obscurin) in C. elegans muscle, J. Mol. Biol. 386 (2009) 976–988.

[10] K.J. Wilson, H. Qadota, P.E. Mains, G.M. Benian, UNC-89 (obscurin) binds to MEL-26, a BTB-domain protein, and affects the function of MEI-1 (katanin) in striated muscle of C. elegans, Mol. Biol. Cell 23 (2012) 2623–2634.

[11] A. Warner, G. Xiong, H. Qadota, T. Rogalski, A.W. Vogl, D.G. Moerman, G.M. Benian, CPNA-1, a copine domain protein, is located at integrin adhesion sites and is required for myofilament stability in Caenorhabditis elegans, Mol. Biol. Cell 24 (2013) 601–616.

[12] H. Qadota, O. Mayans, Y. Matsunaga, J.L. McMurry, K.J. Wilson, G.E. Kwon, R. Stanford, K. Deehan, T.L. Tinley, V.M. Ngwa, G.M. Benian, The SH3 domain of UNC-89 (obscurin) interacts with paramyosin, a coiled-coil protein, in Caenorhabditis elegans muscle, Mol. Biol. Cell 27 (2016) 1606–1620.

[13] H. Qadota, Y. Matsunaga, P. Bagchi, K.I. Lange, K.J. Carrier, W.V. Pols, E. Swartzbaugh, K.J. Wilson, M. Srayko, D.C. Pallas, G.M. Benian, Protein phosphatase 2A is crucial for sarcomere organization in Caenorhabditis elegans striated muscle, Mol. Biol. Cell 29 (2018) 2084–2097.

[14] M.L. Bang, T. Centner, F. Fornoff, A.J. Geach, M. Gotthardt, M. McNabb, C.C. Witt Labeit, C.C. Gregorio, H. Granzier, S. Labeit, The complete gene sequence of titin, expression of an unusual approximately 700-kDa titin isoform, and its interaction with obscurin identify a novel Z-line to I-band linking system, Circ. Res. 89 (2001) 1065–1072.

[15] P. Young, E. Ehler, M. Gautel, Obscurin, a giant sarcomeric Rho guanine nucleotide exchange factor protein involved in sarcomere assembly, J. Cell Biol. 154 (2001) 123–136.

[16] N.A. Perry, M.A. Ackermann, M. Shriver, L.Y. Hu, A. Kontrogianni-Konstantopoulos, Obscurins: unassuming giants enter the spotlight, IUBMB Life 65 (2013) 479–486.

[17] A.L. Bowman, A. Kontrogianni-Konstantopoulos, S.S. Hirsch, S.B. Geisler, H. Gonzalez-Serratos, M.W. Russell, R.J. Bloch, Different obscurin isoforms localize to distinct sites at sarcomeres, FEBS Lett. 581 (2007) 1549–54.

[18] A. Grogan, A. Kontrogianni-Konstantopoulos, Unraveling obscurins in heart disease, Pflugers Arch. 471 (2018) 735–743.

[19] A. Kontrogianni-Konstantopoulos, E.M. Jones, D.B. Van Rossum, R.J. Bloch RJ., Obscurin is a ligand for small ankyrin 1 in skeletal muscle, Mol Biol Cell. 14 (2003) 1138–1148.

[20] P. Bagnato, V. Barone, E. Giacomello, D. Rossi, V. Sorrentino, Binding of an ankyrin-1 isoform to obscurin suggests a molecular link between the sarcoplasmic reticulum and myofibrils in striated muscles, J Cell Biol. 160 (2003) 245–253.

[21] S. Lange, K. Ouyang, G. Meyer, L. Cui, H. Cheng, R.L. Lieber, J. Chen, Obscurin determines the architecture of the longitudinal sarcoplasmic reticulum, J Cell Sci. 122 (2009) 2640–2650.

[22] P.M. Spooner, J. Bonner, A.V. Maricq, G.M. Benian, K.R. Norman, Large isoforms of UNC-89 (obscurin) are required for muscle cell architecture and optimal calcium release in Caenorhabditis elegans, PLoS ONE 7(7)(2012) e40182.

[23] S.B. Geisler, D. Robinson, M. Hauringa, M.O. Raeker, A.B. Borisov, M.V. Westfall, M.W. Russell, Obscurin-like 1, OBSL1, is a novel cytoskeletal protein related to obscurin, Genomics 89 (2007) 521–531.

[24] A. Fukuzawa, S. Lange, M. Holt, A. Vihola, V. Carmignac, A. Ferreiro, B. Udd, M. Gautel M, Interactions with titin and myomesin target obscurin and obscurin-like 1 to the M-band: implications for hereditary myopathies, J. Cell Sci. 121 (2008) 1841–1851.

[25] D. Hanson, P.G. Murray, A. Sud, S.A. Temtamy, M. Aglan, A. Superti-Furga, S.E. Holder, J. Urquhart, E. Hilton, F.D. Manson, P. Scambler, G.C. Black, P.E. Clayton, The primordial growth disorder 3-M syndrome connects ubiquitination to the cytoskeletal adaptor OBSL1, Am. J. Hum. Genet. 84 (2009) 801–806.

[26] P.B. Agrawal, C.R. Pierson, M. Joshi, X. Liu, G. Ravenscroft, B. Moghadaszadeh, T. Talabere, M. Viola, L.C. Swanson, G. Haliloğlu, B. Talim, K.S. Yau, R.J. Allcock, N.G. Laing, M.A. Perrella, A.H. Beggs, SPEG interacts with myotubularin, and its deficiency causes centronuclear myopathy with dilated cardiomyopathy, Am. J. Hum. Genet. 95 (2014) 218–226.

[27] O. Mayans, G.M. Benian, F. Simkovic, D.J. Rigden, Mechanistic and functional diversity in the mechanosensory kinases of the titin-like family, Biochem. Soc. Trans. 41 (2013) 1066–1071.

[28] L.Y. Hu, A. Kontrogianni-Konstantopoulos, The kinase domains of obscurin interact with intercellular adhesion proteins, FASEB J. 27 (2013) 2001–2012.

[29] A.P. Quick, Q. Wang, L.E. Philippen, G. Barreto-Torres, D.Y. Chiang, D. Beavers, G. Wang, M. Khalid, J.O. Reynolds, H.M. Campbell, J. Showell, M.D. McCauley, A. Scholten, X.H. Wehrens, SPEG (Striated Muscle Preferentially Expressed Protein Kinase) Is Essential for Cardiac Function by Regulating Junctional Membrane Complex Activity, Circ. Res. 120 (2016) 110–119.

[30] H. Li, W.A. Linke, A.F. Oberhauser, M. Carrion-Vazquez, J.G. Kerkvliet, H. Lu, P.E. Marszalek, J.M. Fernandez, Reverse engineering of the giant muscle protein titin, Nature 418 (2002) 998–1002.

[31] D.M. Miller, I. Ortiz, G.C. Berliner, H.F. Epstein, Differential localization of two myosins within nematode thick filaments, Cell 34 (1983) 477–490.

[32] H. Qadota, Y. Matsunaga, K.C.Q. Nguyen, A. Mattheyses, D.H. Hall, G.M. Benian, High-resolution imaging of muscle attachment structures in C. elegans, Cytoskeleton (2017) 1–17.

[33] R.H. Waterston, Muscle, In The Nematode Caenorhabditis elegans (1988) (Wood, W.B., ed), pp. 281–335, Cold Spring Harbor Laboratory Press, Cold Spring Harbor, N.Y.

[34] M. Rahman, J.E. Hewitt, F. Van-Bussel, H. Edwards, J. Blawzdziewicz, N.J. Szewczyk, M. Driscoll, S.A. Vanapalli, NemaFlex: a microfluidics-based technology for standardized measurement of muscular strength of C. elegans, Lab on a Chip 18.15 (2018) 2187–2201.

[35] L. Avery, The genetics of feeding in C. elegans, Genetics 133 (1993) 897–917.

[36] D.M. Miller, F.E. Stockdale, J. Karn, Immunological identification of the genes encoding the four myosin heavy chain isoforms of *C. elegans*, Proc. Natl. Acad. Sci. 83 (1986) 2305–2309.

[37] H. Granzier, S. Labeit, Cardiac titin: an adjustable multi-functional spring, J. Physiol. 541(2002) 335–342.

[38] D.G. Moerman, B.D. Williams, Sarcomere assembly in C. elegans muscle, WormBook (2006) 1–16.

[39] K. Gieseler, H. Qadota, G.M. Benian, Development, structure, and maintenance of C. elegans body wall muscle, WormBook (2017) 1–59.

[40] X. Lin, H. Qadota, D.G. Moerman, B.D. Williams, C. elegans PAT-6/actopaxin plays a critical role in the assembly of integrin adhesion complexes in vivo, Curr. Biol. 13(2003) 922–932.

[41] E. von Castelmur, M. Marino, D.I. Svergun, L. Kreplak, Z. Ucurum-Fotiadis, P.V. Konarev, A. Urzhumtsev, D. Labeit, S. Labeit, O. Mayans, A regular pattern of Ig supermotifs defines segmental flexibility as the elastic mechanism of the titin chain, Proc. Natl. Acad. Sci. U S A. 105 (2008) 1186–1191.

[42] I. Agarkova, Perriard J.-C., The M-band: an elastic web that crosslinks thick filaments in the center of the sarcomere, Trends Cell Biol. 15 (2005) 477–485.

[43] S. Lange, N. Pinotsis, I. Agarkova, E. Ehler, The M-band: The underestimated part of the sarcomere, BBA-Mol. Cell Res. 1867 (2020) 118440.

[44] F. Berkemeier, M. Bertz, S. Xiao, N. Pinotsis, M. Wilmanns, F. Grater, M. Rief, Fast-folding alpha-helices as reversible strain absorbers in the muscle protein myomesin, Proc. Natl. Acad. Sci. USA 108 (2011) 14139–14144.

[45] S. Pernigo, A. Fukuzawa, A.E.M. Beedle, M. Holt, A. Round, A. Pandini, S. Garcia-Manyes, M. Gautel, R.A. Steiner, Binding of myomesin to obscurin-like-1 at the muscle M-band provides a strategy for isoform-specific mechanical protection, Structure 25 (2017) 107–120.

[46] P.J. Bujalowski, A. F. Oberhauser, Tracking unfolding and refolding reactions of single proteins using atomic force microscopy methods, Methods 60 (2013) 151–160.

[47] M. Rabbi, P.E. Marszalek, Measuring Protein Mechanics by Atomic Force Microscopy

[48] M.D. Hossain, S. Furuike, K. Kinosita Jr, The rotor tip inside a bearing of a thermophilic F1-ATPase is dispensable for torque generation, Biophys. J. 90 (2006) 4195–4203.

[49] E.-L. Florin, M. Rief, H. Lehmann, M. Ludwig, C. Dornmair, V.T. Moy, H.E. Gaub, Sensing specific molecular interactions with the atomic force microscope. Biosens. and Bioelectr. 10 (1995) 895–901.

[50] M. Kim, K. Abdi, P.E. Marszalek, Fast and forceful refolding of stretched alpha-helical solenoid proteins, Biophys. J. 98 (2010) 3086–3092.

[51] C. Bustamante, J.F. Marko, S. Smith S, Entropic elasticity of lambda-phage DNA. Science 265 (1994) 1599–1600.

[52] J.F. Marko, E.D. Siggia, Stretching DNA, Macromolecules 28 (1995) 8759–8770.

[53] E. Hannak, K. Oegema, M. Kirkham, P. Gonczy, B. Habermann, A.A. Hyman, The kinetically dominant assembly pathway for centrosomal asters in Caenorhabditis elegans is gamma-tubulin dependent, J. Cell Biol. 157 (2002) 591–602.

[54] M.C. Hresko, B.D. Williams, R.H. Waterston, Assembly of body wall muscle and muscle cell attachment structures in Caenorhabditis elegans, J. Cell Biol. 124 (1994) 491–506.

[55] M.L. Nonet, K. Grundahl, B.J. Meyer, J.B. Rand, Synaptic function is impaired but not eliminated in C. elegans mutants lacking synaptotagmin, Cell 73 (1993) 1291–1305.

[56] K.J. Wilson, H. Qadota, G.M. Benian, Immunofluorescent localization of proteins in Caenorhabditis elegans muscle, Methods Mol. Biol. 798 (2012) 171–181.

[57] H. Qadota, K.B. Mercer, R.K. Miller, K. Kaibuchi, G.M. Benian, Two LIM domain proteins and UNC-96 link UNC-97/pinch to myosin thick filaments in Caenorhabditis elegans muscle, Mol. Biol. Cell 18 (2007) 4317–4326.

[58] R. Francis, R.H. Waterston, Muscle cell attachment in Caenorhabditis elegans, J. Cell Biol. 114 (1991) 465–479.

[59] C. Nussbaum-Krammer, M.F. Neto, R.M. Brielmann, J.S. Pedersen, R.I. Morimoto, Investigating the spreading of toxicity of prion-like proteins using the metazoan model organism C. elegans, J. Vis. Exp. 95 (2015) 52321.

[60] J.E. Hewitt, A.K. Pollard, L. Lesanpezeshki, C.S. Deane, C.J. Gaffney, T. Etheridge, N.J. Szewczyk, S.A. Vanapalli, Muscle strength deficiency and mitochondrial dysfunction in a muscular dystrophy model of Caenorhabditis elegans and its functional response to drugs, Disease Mod. & Mechan. 11.12 (2018).

